# Assessing the response of small RNA populations to allopolyploidy using resynthesized *Brassica napus* allotetraploids

**DOI:** 10.1101/441022

**Authors:** Paulina Martinez Palacios, Marie-Pierre Jacquemot, Marion Tapie, Agnès Rousselet, Mamoudou Diop, Carine Remoue, Matthieu Falque, Andrew Lloyd, Eric Jenczewski, Gilles Lassalle, Anne-Marie Chevre, Christine Lelandais, Martin Crespi, Philippe Brabant, Johann Joets, Karine Alix

## Abstract

Allopolyploidy, combining interspecific hybridization with whole genome duplication, has had significant impact on plant evolution. Its evolutionary success is related to the rapid and profound genome reorganizations that allow neo-allopolyploids to form and adapt. Nevertheless, how neo-allopolyploid genomes adapt to regulate their expression remains poorly understood. The hypothesis of a major role for small non-coding RNAs (sRNAs) in mediating the transcriptional response of neo-allopolyploid genomes has progressively emerged. Generally, 21-nt sRNAs mediate post-transcriptional gene silencing (PTGS) by mRNA cleavage whereas 24-nt sRNAs repress transcription (transcriptional gene silencing, TGS) through epigenetic modifications. Here, we characterize the global response of sRNAs to allopolyploidy in *Brassica*, using three independently resynthesized *B. napus* allotetraploids surveyed at two different generations in comparison with their diploid progenitors. Our results suggest an immediate but transient response of specific sRNA populations to allopolyploidy. These sRNA populations mainly target non-coding components of the genome but also target the transcriptional regulation of genes involved in response to stresses and in metabolism; this suggests a broad role in adapting to allopolyploidy. We finally identify the early accumulation of both 21- and 24-nt sRNAs involved in regulating the same targets, supporting a PTGS-to-TGS shift at the first stages of the neo-allopolyploid formation. We propose that reorganization of sRNA production is an early response to allopolyploidy in order to control the transcriptional reactivation of various non-coding elements and stress-related genes, thus ensuring genome stability during the first steps of neo-allopolyploid formation.

## Introduction

Through the course of their evolution and environmental adaptation, living organisms repeatedly face threats of diverse nature, against which many defense responses are initiated by the genome itself. This was first established by Barbara McClintock who considered interspecific hybridization a “specific shock that forced the genome to restructure itself in order to overcome a threat to its survival” (1984). Since then, concerted efforts have been made by the scientific community to decipher the modalities of such genome reorganization in the context of the formation of new species.

Work in this area has notably focused on the consequences of interspecific hybridization and allopolyploidy in plants. Allopolyploidy combines interspecific hybridization with whole genome duplication; this is a prominent force in the evolution of many eukaryote lineages (reviewed by e.g. Soltis and Soltis 2012, Chen and Birchler 2013) including plants where it has been particularly pervasive (Jiao et al. 2011, Alix et al. 2017). It has been shown that neo-allopolyploid genomes undergo numerous changes, at both the structural and functional levels. These rapid and profound genome reorganizations (Madlung and Wendel, 2013) explain the ability of allopolyploids to compete with their parental species and to colonize new habitats (Hegarty and Hiscock 2008), as well as the large extent of their adaptability and thus evolutionary success. However, in-depth analysis of the molecular mechanisms underpinning the regulation of genome expression during allopolyploidization has rarely been undertaken, except in the model plant Arabidopsis (Shi et al. 2015 and references therein). Through these few studies, the hypothesis of a major role for small RNAs in mediating the expression of neo-allopolyploid genomes to genomic shock has progressively emerged.

Plant small non-coding RNAs have been subjected to various classifications (e.g. Axtell 2013; Martínez de Alba et al. 2013). Based on their modes of biogenesis, sRNAs may be divided into two major classes (Martínez de Alba et al. 2013; Mirouze 2012): (i) microRNAs (miRNAs), generally 21 nucleotides (nt) in length, which are produced from single-stranded hairpin RNAs transcribed from endogenous *MIR* genes (Voinnet 2009), and (ii) small (or short) interfering RNAs (siRNAs) that represent a highly diverse class of 21- to 24-nt sRNAs (and thus subject to numerous sub-classifications). These siRNAs are all produced from double-stranded RNA (dsRNA) precursors of various origins, such as inverted repeats, natural antisense overlapping transcripts, or dsRNAs synthesized from single-stranded RNAs (ssRNAs) by RNA-dependent RNA polymerases (RDRs). In this latter class, the primary ssRNAs result from the transcription of transposable elements (TEs) or other repetitive sequences (Bond and Baulcombe 2014). The various types of sRNAs have different modes of action; miRNAs and 21-nt siRNAs have been demonstrated to mostly mediate post-transcriptional gene silencing (PTGS) by endonucleolytic cleavage of messenger RNA targets or repression of translation, while 24-nt siRNAs usually repress gene activity by transcriptional gene silencing (TGS) through modifications of epigenetic chromatin marks (DNA methylation, histone modifications) at specific genomic loci. For every sRNA, the highly specific action of gene repression is based on base-pairing (Mallory and Vaucheret 2006), with selectivity being mediated by sequence similarity to the sRNA which has been loaded into an Argonaute (AGO) protein – the major component of the effector silencing protein complex RISC (RNA-induced silencing complex).

Few studies have been dedicated to the analysis of sRNA mobilization during interspecific hybridization (tomato, Shivaprasad et al. 2012) or allopolyploidization (Arabidopsis, Ha et al. 2009; wheat, Kenan-Eichler et al. 2011), but those that have been undertaken demonstrate significant changes in sRNA expression profiles accompanying the formation of hybrid/allopolyploid genomes. As these studies analysed the mobilization of sRNAs by pooling individual samples to represent each generation, the stochasticity or not of these responses has remained unclear. Addressing this question by evaluating the global dynamics of sRNA populations to independent allopolyploidization events, across several generations, has therefore remained a major important task. This was our aim in this study. Using the *Brassica* model, we undertook high-throughput sequencing of small RNAs, characterizing global changes in sRNA accumulation during the formation of three independent resynthesized *B. napus* allotetraploids in comparison with their progenitors.

*Brassica* represents a good model to study speciation by allopolyploidy, with the allotetraploid species *B. napus* (oilseed rape, AACC, 2n=38) originating from interspecific hybridization between the diploid species *B. rapa* (AA, 2n=20) and *B. oleracea* (CC, 2n=18) less than 10 000 years ago. As the current diploid *Brassica* species are still closely related to the true diploid progenitors of *B. napus*, the allotetraploid can be resynthesized artificially, and therefore represents an excellent model for analysis of the genomic consequences of polyploidy over different timescales (Jenczewski et al. 2013). In the present study, we characterized the global response of sRNA populations to allopolyploidy in *Brassica*, by comparing the sRNA distributions of two generations (S1, S5) of three resynthesized *B. napus* allotetraploids (originating from the same diploid genotypes) to those of their diploid progenitors (fig. 1). Using high-throughput sequencing of sRNA populations, we analysed the size distributions across these lines, and annotated the sRNA fractions showing differential abundances across the generations under survey, combining bioinformatic data with experimental molecular validation. For specific sRNA populations, the dynamics of accumulation was detailed using additional allotetraploid generations (S0, S3) as well as the first hybrids (F1), by quantitative RT-PCR (details of the original plant material analysed is provided in fig. 1). Here, we report the global dynamics of sRNA accumulation from the diploid to the allotetraploid genomic context, emphasizing the results that identified sRNA populations responding to the “genomic stress” triggered by allopolyploidy. We discuss the mechanisms of action of small non-coding RNAs established during allopolyploidization, and the putative role of sRNAs in the success of neo-allopolyploids.

**Figure 1.**
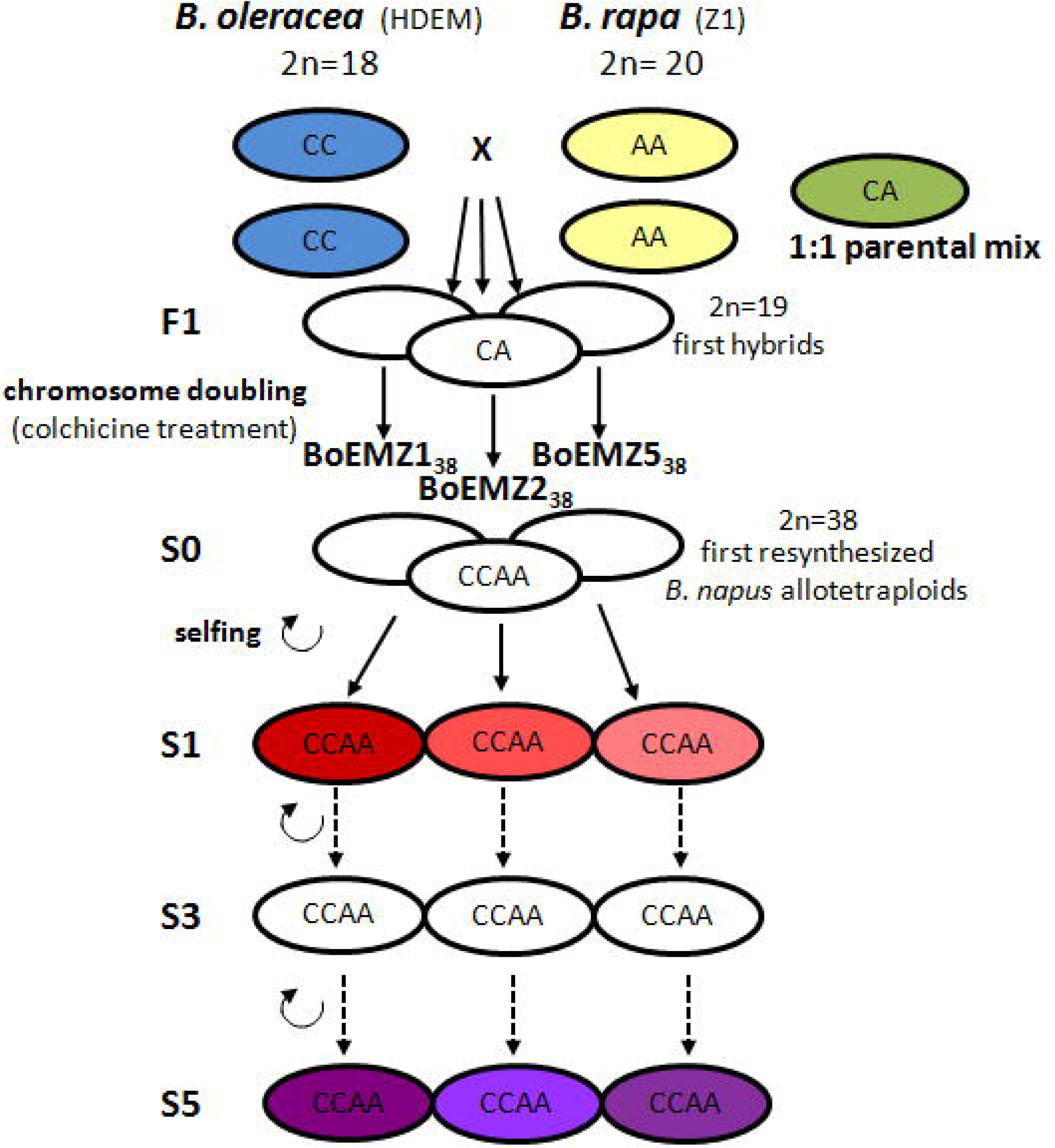
Pedigree of the three resynthesized *B. napus* allotetraploids analysed in the present study, and identification of the genotypes used for RNA extraction and subsequent construction of small RNA libraries and Illumina sequencing. A total of eleven small RNA libraries were constructed and sequenced, corresponding to RNA extracts from the diploid progenitors (with two biological replicates for each progenitor; with *B. oleracea* in blue and *B. rapa* in yellow) as well as a 1:1 parental RNA mix used as a technical replicate (in green), and RNA extracts from the three resynthesized *B. napus* allotetraploids surveyed at two different generations of selfing, S1 (in red) and S5 (in purple). The F1 hybrids and the allotetraploid generations S0 and S3 (not colored) were used to detail the dynamics of accumulation of specific sRNA populations by qRT-PCR.

## Results

To study the global impact of allopolyploidy on small RNA populations, we analyzed a total of ~15,000,000 Illumina sequence reads of 18-26-nt in size (bioinformatic treatment, normalization of the read counts, and further analyses of the sequencing data are detailed in materials and methods). These reads were generated from sRNA-enriched libraries constructed from three resynthesized *B. napus* allotetraploids at two different generations of selfing and their diploid progenitors *B. oleracea* and *B. rapa* (fig. 1). To allow integration of the current data with available functional data generated from the same plant material (Albertin et al. 2006, 2007) we specifically collected and used stem tissues.

### Expression levels of small RNAs experienced rapid changes following allopolyploidy

Size distribution of small RNAs (normalized in reads per million, RPM, to allow comparisons between libraries with different sequencing depths) showed that the 24-nt class was the most abundant class in all libraries, accounting on average for 46.54 % of the total number of reads per library, followed by the 21-nt class that represented on average 20.63 % of the reads per library (supplementary table S1, Supplementary Material on line). Interestingly, the 21-nt sRNAs were significantly more abundant in the S1 generation than in both the S5 generation (*P*<0.05) and the parental generation by comparison to the mid-parent value (MPV that represents the additive hypothesis from the parental generation; *P*<0.005). By contrast, no significant differences were observed when comparing the abundance of the 21-nt sRNAs in the S5 generation to the MPV (*P*=0.22) (fig. 2*A*). At the S1 generation, the increase of the 21-nt sRNA class was accompanied by a significant decrease of the 24-nt class by comparison to the MPV (*P*<0.05). In the same manner, the 24-nt class was significantly less abundant in the S1 than in the S5 generation (*P*<0.01). As for the 21-nt class, no significant differences were observed between the relative abundances of the 24-nt sRNAs in the S5 generation and the parental generation (tested still using the MPV, *P*=0.22).

**Figure 2.**
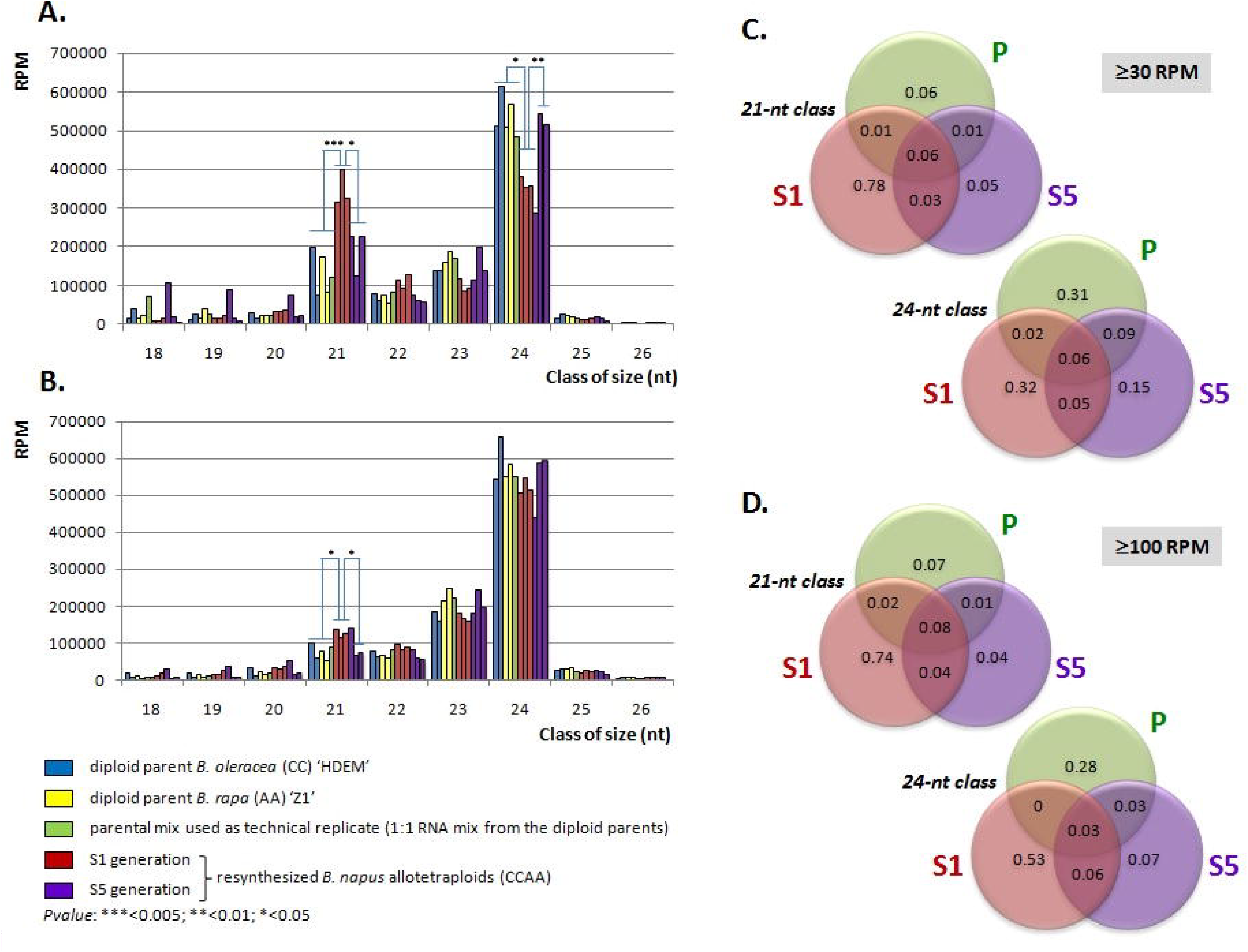
Size distribution of small RNAs in three resynthesized *B. napus* allotetraploids (BoEMZ1_38_, BoEMZ2_38_, BoEMZ5_38_ – in order) surveyed at two different generations of selfing (S1 and S5) in comparison with their diploid progenitors *B. oleracea* (HDEM) and *B. rapa* (Z1). (**A**) Size distribution of redundant sRNAs is provided with (**B**) size distribution of unique reads (illustrating the diversity of sRNAs that are different in nucleotide sequences). (**C,D**) The diversity for sRNAs with expression levels ≥30 RPM (**C**) and ≥100 RPM (**D**) was analyzed across the generations, for both the 21-nt and the 24-nt classes; the results indicated that for each condition of expression, 78%-74% and 32%-53% of the 21-and 24-nt reads analyzed respectively, were S1-specific, suggesting the expression of specific sRNA populations in response to allopolyploidy. Only 6 to 17% of the sRNAs surveyed were common to the diploid progenitors and the resynthesized *B. napus* allotetraploids (making there no distinction between S1 and S5). Venn diagrams show fractions of total 21- and 24-nt unique reads with expression levels ≥30 RPM and ≥100 RPM for each generation, with sequencing data obtained from each library being pooled per generation: diploid parents ‘P’ (pool of 5 libraries), S1 generation ‘S1’ (3), S5 generation ‘S5’ (3). Details of quantifications are provided in tables S1 and S2 (supplementary material.

We tested if the variations observed for the 21-nt and 24-nt classes across the generations were mainly (i) quantitative, with the same populations of sRNAs showing different levels of expression across the generations, or (ii) qualitative, with populations of sRNAs being expressed specifically in each generation. To do so, we determined the composition of the sRNA diversity for the 21-nt and 24-nt classes. After removing sequence redundancy, the 24-nt class was the most diverse in all the libraries (fig. 2*B*), and accounted on average for 55.36% of the unique reads per library (supplementary table S2, Supplementary Material on line). The abundance of the 24-nt class was related to the presence of numerous unique 24-nt sRNAs, which were all expressed on average at relatively low levels (characteristic of the 24-nt class already shown in previous studies, e.g. Lelandais-Brière et al. 2009). In contrast, we observed a drastic reduction in the 21-nt class after removing read redundancy (accounting, on average, for only 9.41% of the unique reads per library), suggesting high levels of accumulation for less diverse 21-nt sRNAs in the three generations surveyed (fig. 2*B*). The only significant differences we detected across the generations when comparing the diversity (i.e. the number of unique sRNA reads) among the 21-nt class or the 24-nt class corresponded to a slight increase in the level of the 21-nt sRNA diversity estimated for the S1 generation in comparison to the ones estimated for the parental generation (*P*<0.05 against the MPV) and the S5 generation (*P*<0.05). We repeated this analysis considering only sRNAs with moderate (RPM ≥ 30) or high (RPM ≥ 100) expression to limit the impact of stochastic background sRNA expression on the results (Fahlgren et al. 2009). In these conditions, the results highlighted a significantly higher diversity within the S1 generation, indicating that abundant sRNAs specifically accumulate in the S1 generation of the resynthesized *B. napus* allotetraploids (fig. 2*C* and 2*D*). It can also be noted that only a few sRNAs showed high accumulation levels in both the diploid progenitors and at least one generation of the resynthesized *B. napus* allotetraploids. Accordingly, the qualitative analysis of sRNA diversity across the three surveyed generations demonstrated the immediate mobilization of specific sRNA populations in response to allopolyploidy.

Altogether, the results obtained from the analysis of the size-specific distribution of small RNAs across the three generations surveyed, clearly showed that early generations (S1) of the *B. napus* neo-allopolyploids exhibited a strong over-expression of 21-nt sRNAs, with specific sRNA populations accumulating at high levels in the S1 only, accompanied by a slight under-expression of 24-nt sRNAs in comparison to the MPV. We thus carried out detailed bioinformatic annotation of the two classes of reads, to identify the sRNA populations responsible for the deviation in sRNA accumulation patterns observed during the formation of the *Brassica* neo-allotetraploids.

### The global expression of microRNAs was not significantly affected by allopolyploidy

We first considered the subset of small RNAs corresponding to microRNAs. We identified conserved microRNAs (miRNAs) by comparing the entire set of unique reads for the 21-nt and 24-nt sRNA classes to the plant miRBase database (release 19). No conserved miRNAs were identified among the 24-nt reads in accordance with the rarity of 24-nt miRNAs in the plant miRBase. Among the 21-nt class, more than one third of the sRNA reads represented previously known microRNAs (fig. 3*A*). In total, we retrieved 133 unique known miRNAs corresponding to 28 different microRNA families (supplementary table S3, Supplementary Material on line) according to the classification published in miRBase (Kozomara and Griffiths-Jones 2014); these included 24 different miRNA families that had been previously identified in *Brassica*. In terms of diversity, most of the retrieved miRNA families with expression levels ≥ 30 RPM were found in the three generations analysed (76%), with no known miRNAs displaying specific accumulation in the S1 generation. It can be noticed that few miRNAs accumulated specifically in the diploid parents (14% - miR398, miR400, miR845) or the S5 generation (5% - miR399), or in both the parents and the S5 generation (5% - miR824) (fig. 3*B* and details in supplementary table S3, Supplementary Material on line).

**Figure 3.**
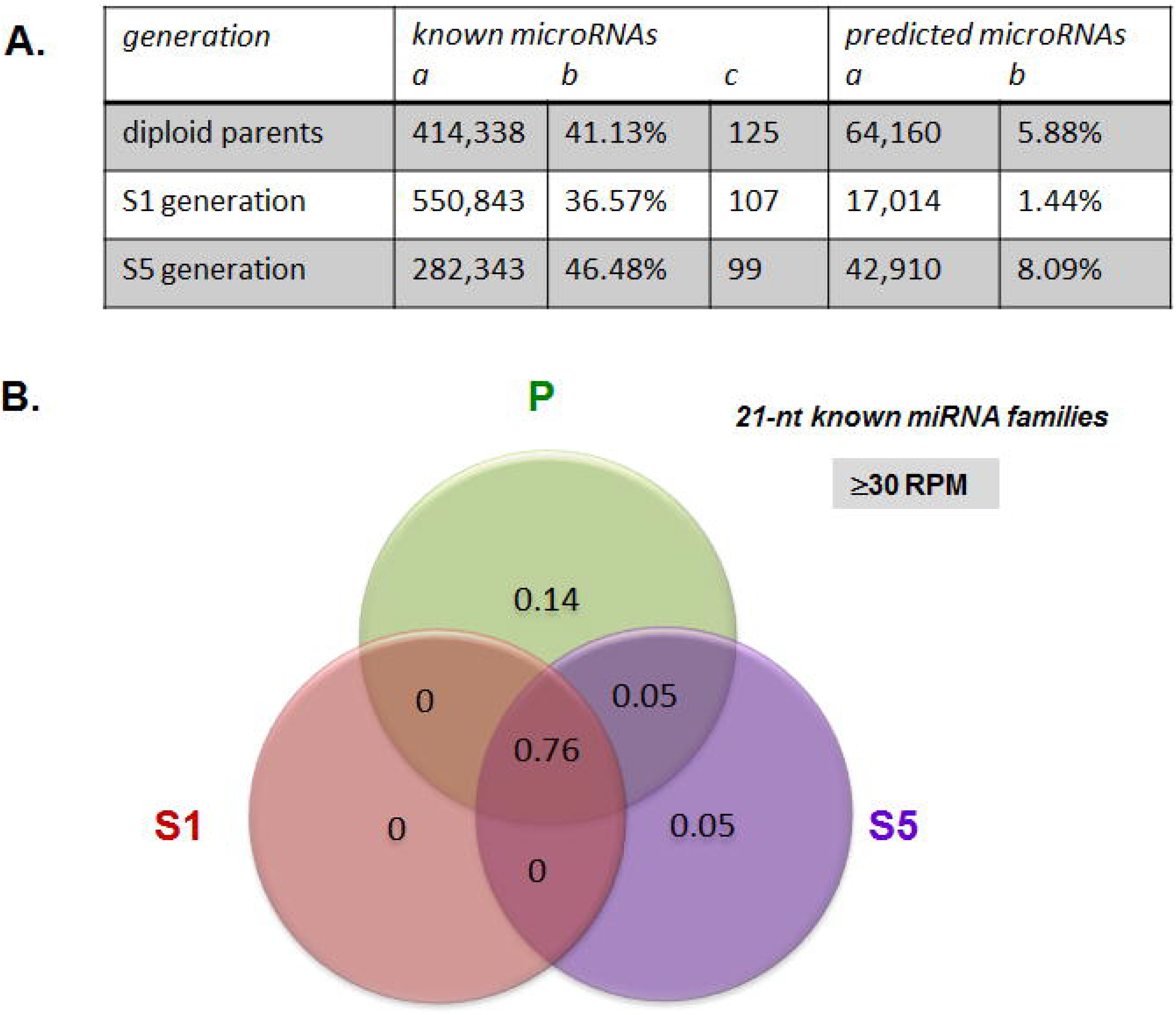
Characterization of the sRNAs annotated as microRNAs in the resynthesized *B. napus* allotetraploids surveyed at the two generations S1 and S5 in comparison with their diploid parents. (**A**) Summary table reporting the number of raw read counts (column a), the percentage of the 21-nt class (b) and the number of unique reads (c) corresponding to known or predicted microRNAs identified among the 21-nt reads. (**B**) Venn diagram showing the distribution of the 28 known microRNA families (according to miRBase v.19) that were present in the sequencing data with expression levels ≥30 RPM. Most of the microRNA families were found in the three different generations (76%); only 5 miRNA families were more abundant in the parental (14%) or the S5 generation (5%), or in both the parents and the S5 lines (5%). For this analysis, the sequencing data obtained from each library were pooled by generation: diploid parents ‘P’ (pool of 5 libraries), S1 generation ‘S1’ (3), S5 generation ‘S5’ (3). Distribution of known miRNAs appears thus as the opposite to the one depicted for the whole 21-nt class and reported in fig. 2*C*.

To complete our analysis, we performed microRNA prediction on the non-annotated 21-nt and 24-nt fractions of the libraries with miRCat from the UEA plant sRNA toolkit (http://srna-tools.cmp.uea.ac.uk/). This toolkit identifies mature miRNAs as well as their precursors (Moxon et al. 2008). No miRNAs were predicted among the 24-nt reads, while a total of 274 unique reads were predicted as miRNA candidates from the 21-nt class according to hairpin prediction (fig. 3*A*). The full set of the putative *MIR* genes, characterization of the corresponding miRNA precursors, and locations of the genes along the *B. rapa* genome are given in supplementary table S4 (Supplementary Material on line). The identification of the miRNA* strands in addition to the guide miRNA strands for 92 out of the collection of 274 newly predicted *Brassica* microRNAs adds robustness to this *Brassica* miRNA prediction that may be useful for further investigation.

For the known miRNAs retrieved from our sRNA sequencing data, we observed no quantitative differences when comparing their global relative abundance across the different generations surveyed (ANOVA, *P*=0.72). Similarly, no significant differences were observed when considering the relative abundance of the whole fraction of sRNAs annotated as miRNAs (known and predicted) between the three generations (ANOVA, *P*=0.71). Nevertheless, analyzing in detail the accumulation levels of specific miRNAs suggested that, while global miRNA abundance is stable, some individual miRNAs displayed some differential accumulation levels across the generations (supplementary table S3, Supplementary Material on line). We performed quantification of expression by real-time quantitative RT-PCR (qRT-PCR) of a set of four known microRNAs (miR159a, miR160a, miR166a, miR168a). Stable accumulation across the generations was validated for three microRNAs miR159a, miR160a and miR166a in agreement with the sequencing data; these were further used as references (supplementary figure S1, Supplementary Material on line). By contrast, the relative abundance of miR168 was significantly different across the generations tested (ANOVA, *P*=0.0001232) and was notably over-accumulated in the S5 generation compared to the S1 generation (*P*<0.01) in accordance with the sequencing data (4800 vs. 1900 RPM).

In summary, our bioinformatic analysis of the dynamics of the expression of miRNAs following allopolyploidy indicated that the microRNA fraction was not responsible for the immediate and drastic changes we observed in the S1 generation.

### Non-additive small RNAs targeting protein-coding genes were mainly associated with functions related to response to stresses or metabolism

We used a set of 263 proteins for which their immediate response to allopolyploidy (for changes in their abundance) had been previously analyzed: 111 (42.2%) had additive expression and 152 (57.8%) were identified as non-additive (i.e. with abundances that differ from the mid-parent values) (Albertin et al. 2007). We retrieved all the *Brassica* homologous genes from the two diploid species *B. rapa* and *B. oleracea* to constitute ‘*Brassica* functional group sequences’ (one group per protein) onto which we mapped all the 21-and 24-nt small RNA populations, for each generation separately (parents, S1, S5). From the 263 proteins surveyed, a total of 180 proteins corresponding to 64 additive proteins and 116 non-additive proteins had their coding genes targeted by small RNAs (both 21-and 24-nt). Interestingly, these results indicate that small RNAs preferentially targeted proteins identified as non-addititive in response to allopolyploidy (c^2^ test for homogeneity: χ^2^=10.39, *P*=0.001265). On the other hand, we observed a significant over-representation of additive proteins among those that have both their cDNA and genomic sequences targeted by sRNAs (fig. 4*A*; χ^2^=11.73, *P*=0.000615). Forty-four proteins (out of the 180; ~25%) were targeted by populations of sRNAs that were non-additively accumulated in the neo-allopolyploids compared to their diploid progenitors (i.e. against MPV, *P*<0.05). As shown in figure 4*B*, the genes encoding these proteins were regulated mainly at the genomic level only (29/44, 65.9%) either by 21-nt small RNAs (13/44, 29.5%) or by 24-nt small RNAs (28/44, 63.6%). For this set of 44 proteins targeted by non-additive sRNAs, a third (14/44) had additive expression while two-thirds (30/44) had non-additive expression. This proportion was not different to that observed for proteins targeted by additive sRNAs (χ^2^=0.51, *P*=0.4728) and thus the additivity of sRNAs had no discernible influence on the additivity of protein expression.

**Figure 4.**
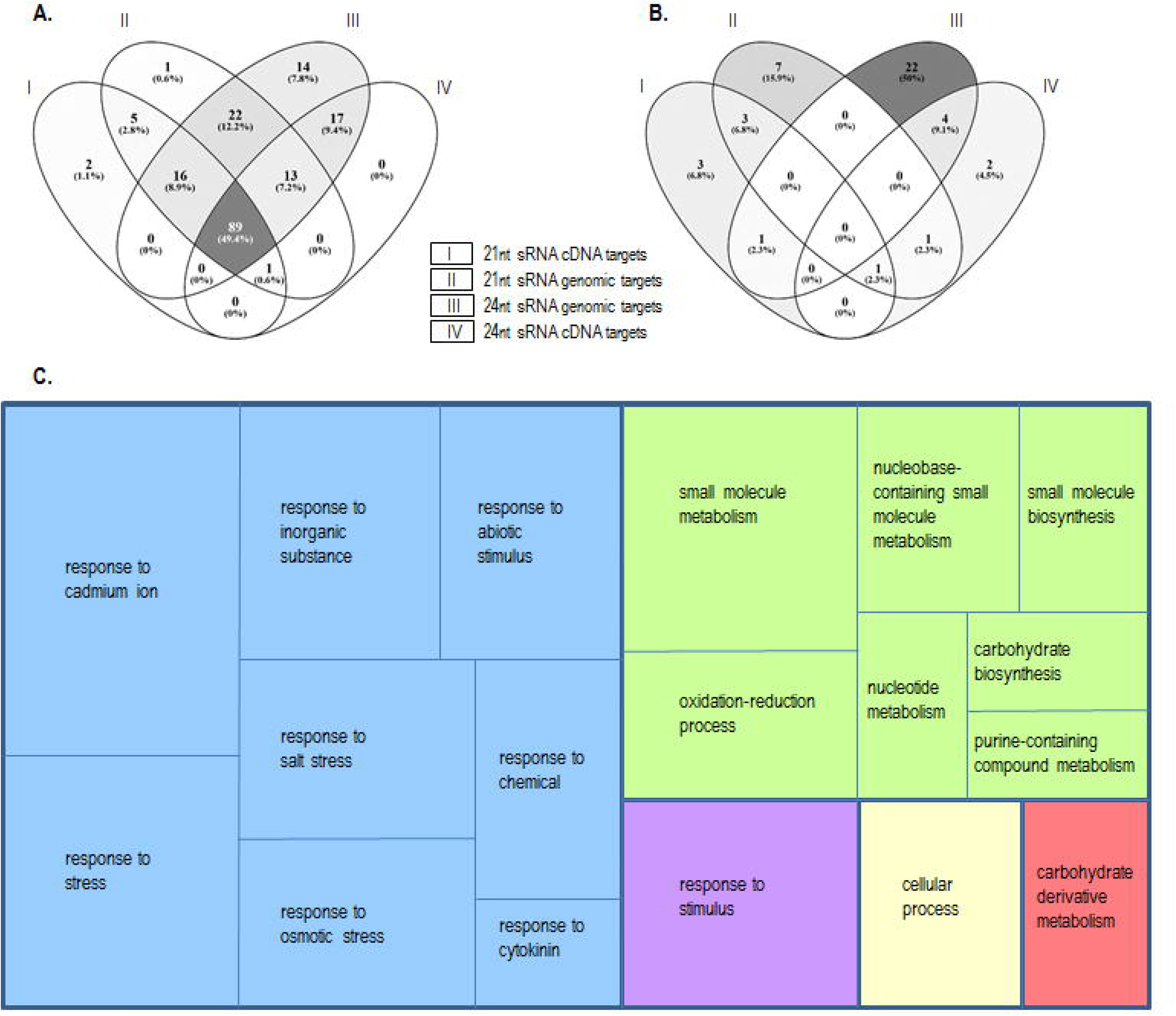
Regulatory relationships between sRNAs and protein expression from the analysis of a set of 263 proteins that were previously analysed for their changes following allopolyploidy. (**A**) A total of 180 proteins were identified as putative targets of the sRNAs present in our sequencing data, with 89 of them that might be regulated at both the transcript and genomic levels by populations of 21- and 24-nt sRNAs. (**B**) From these 180 proteins, 44 proteins including 30 non-additive proteins (Albertin et al. 2007) were targeted by non-additively accumulated sRNAs in the neo-allopolyploids compared to the diploid progenitors, mainly at the genomic level, indicating a TGS control. (**C**) GO term enrichment for biological processes led to the identification of functions associated mainly with responses to stresses and stimuli (blue-violet) or metabolism (green-red). Venn diagrams were drawn using Venny (http://bioinfogp.cnb.csic.es/tools/venny/).

We looked for enriched GO terms for biological processes within the 44 proteins targeted by differentially accumulated sRNAs in response to allopolyploidy. We obtained 18 significant GO terms with the majority of them being associated with responses to various stresses and stimuli while others were related to metabolism (fig. 4*C*). Notably, among the 29 genes involved in response to stresses and stimuli, 24 had reduced accumulation of their corresponding sRNA populations in the resynthesized *Brassica* allotetraploids in comparison to their diploid parents (*P*<0.05 against the MPV including a few with *P*<0.01 or <0.005). According to previous proteomic results, most of the corresponding proteins displayed up-regulated or additive patterns in the S1 neo-allotetraploids (Albertin et al. 2007). This is in accordance with the dynamics of the sRNAs we depicted here; a decrease in the level of sRNAs targeting these stress related genes/mRNAs ended in over-expression or at least maintenance of the expression of the encoded proteins. Similar expression patterns were observed for the 9 genes involved in carbohydrate metabolism and other metabolism processes, with 8 of them corresponding to under-accumulated sRNA populations in the S1 neo-allopolyploids compared to the diploid progenitors (*P*<0.01 against the MPV, with few cases with *P*<0.005 or <0.001) while 6 of these proteins were depicted as up-regulated or additive. Interestingly, the non-additive sRNA populations regulated the expression of their gene targets preferentially at the genomic level, suggesting a predominant TGS control.

The results obtained from the analysis of sRNAs targeting protein-coding genes clearly demonstrated that non-additively accumulated proteins were the preferential targets of sRNA-based regulation. The changes we encountered corresponded mainly to a decrease in the sRNA amounts within the 24-nt class, such changes being consistent with the reduced 24-nt accumulation in S1 described above. To explain the overall increase of the 21-nt class, we then focused our analysis on the sRNAs potentially derived from transposable elements and non-coding intergenic DNA.

### The expression changes of small RNAs derived from transposable elements in response to allopolyploidy were TE-dependent

For each of the two 21-nt and 24-nt classes, we identified small RNAs derived from transposable elements (thereafter denominated siRNAs for small interfering RNAs, Axtell et al. 2013) by subjecting the sequence reads to BLAST analyses using Repbase in addition to a locally developed *Brassica* transposable element database (see materials and methods), first with no distinction between TE families. Among the 21-nt class, relatively few reads were similar to TE sequences corresponding on average to 7.09% (representing 71,454 raw read counts), 3.47% (52,207) and 6.80% (41,299) of the 21-nt reads from the parental, the S1 and the S5 generations, respectively. No significant quantitative differences could be found when comparing the relative abundance of the 21-nt fraction annotated as TE-derived siRNAs between the three different generations (ANOVA, *P*=0.61). Within the 24-nt class, TE-siRNAs were estimated to represent on average 15.71% (597,714 raw read counts), 12.37% (189,912) and 13.92% (236,162) of the 24-nt reads from the parental, S1 and S5 generations, respectively. The relative abundance of the 24-nt TE-derived siRNAs was weakly but significantly different between the generations (ANOVA, *P*=0.03), with a significant decrease in the abundance of the 24-nt TE-siRNAs in the S1 generation in comparison to the MPV (*P*<0.05) and S5 generation (*P*<0.01). No significant quantitative differences in 24-nt TE-siRNA abundance were observed between the S5 generation and the MPV (*P*=0.28). The fraction of sRNAs annotated as TE-siRNAs thus contributed, at least partly, to the overall decrease in the 24-nt class observed in S1 (fig. 2*A*).

We next annotated the TE-siRNA fraction, for both the 21-nt and 24-nt classes, to describe in detail the changes observed in the TE-siRNA populations. Among the 21-nt class, only expression levels of siRNAs derived from LTR-retrotransposons differed significantly between the libraries (ANOVA, *P*=0.025): their abundance was significantly higher in the two generations S1 and S5 of the neo-allotetraploids in comparison to the MPV (*P*<0.01 in both comparisons, fig. 5*A*), while no significant difference was found between the generations S1 and S5 (*P*=0.9). We then annotated siRNAs according to their origins from *Gypsy* or *Copia* LTR-retrotransposons (fig. 5*B*); while the 21-nt *Copia*-siRNAs were additively expressed in the neo-allotetraploids in comparison to the MPV (ANOVA, *P*=0.2), the 21-nt *Gypsy*-siRNAs were significantly over-represented in the S1 and S5 generations in comparison to the expected MPV (*P*<0.05 and *P*<0.01, respectively). When analyzing the 24-nt TE-siRNA fraction, the most significant changes observed in the expression levels of siRNAs (fig. 5*A*, 5*C*) were related to (i) the relative increase in siRNAs derived from LTR-retrotransposons – notably from *Gypsy* when comparing the S5 generation to MPV and S1 (*P*<0.01 and *P*<0.005, respectively) and (ii) the highly significant decrease in other TE-siRNAs in the S1 generation compared to MPV (*P*<0.005 for siRNAs derived from LINEs and *P*<0.05 for siRNAs derived from SINEs or class 2 TIR-transposons). The expression dynamics of 24-nt TE-siRNAs during the formation of the neo-allotetraploids were independently validated by qRT-PCR (supplementary fig. S2, Supplementary Material on line).

**Figure 5.**
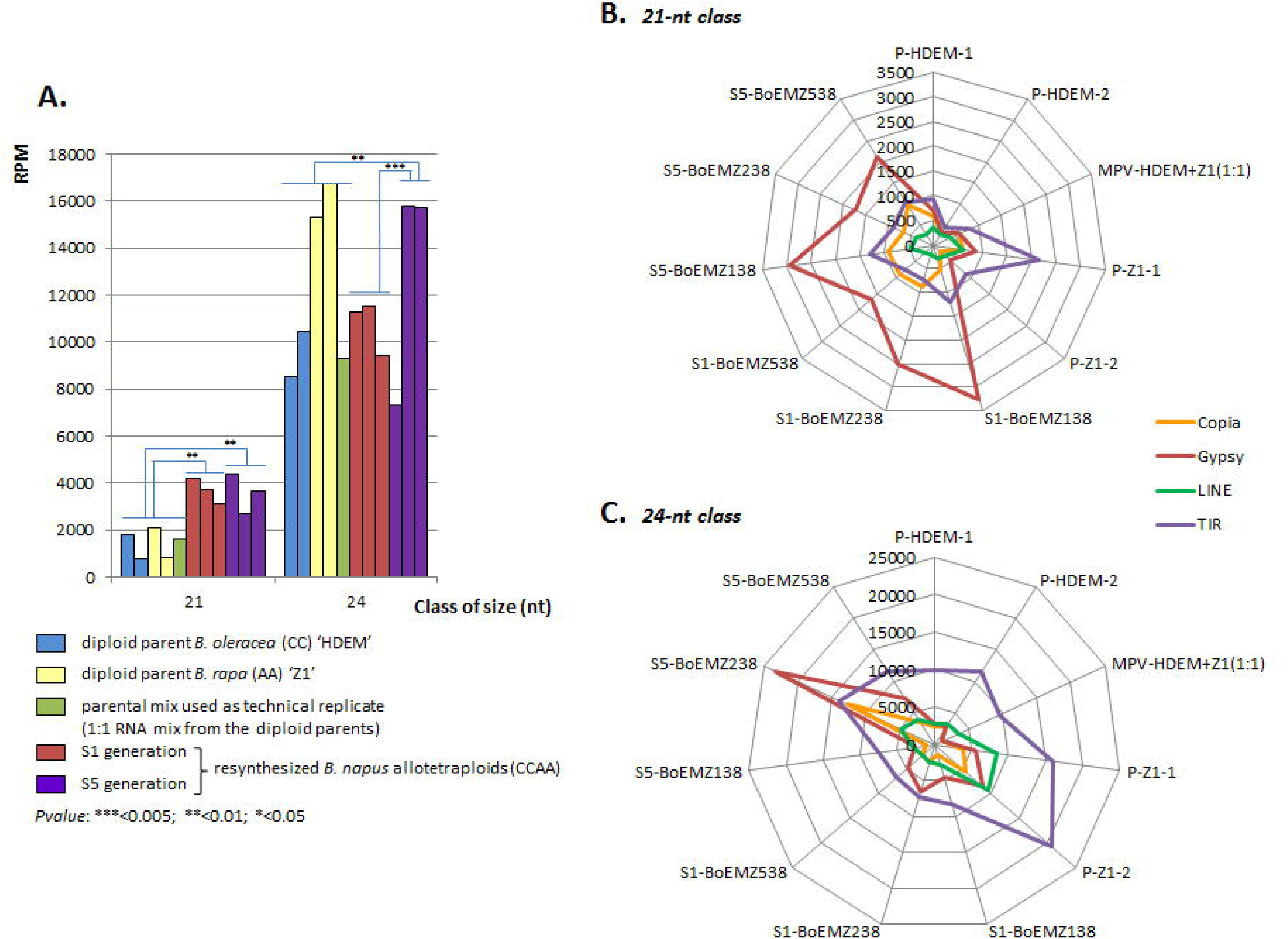
Differential expression of 21-nt and 24-nt siRNAs derived from transposable element sequences in the three resynthesized *B. napus* allotetraploids surveyed at the two generations S1 and S5 in comparison with their diploid progenitors. (**A**) The normalized abundances (in reads per million, RPM) of siRNAs derived from LTR-retrotransposons were estimated for each of the 21-nt and 24-nt classes of sRNAs. We observed a significant increase of the 21-nt siRNAs in the S1 generation in comparison to the parental generation, which was maintained in the S5 generation. The 24-nt siRNAs displayed a significant increase only in the S5 generation in comparison to the parental and the S1 generations, which was particularly marked for the BoEMZ2_38_ and BoEMZ5_38_ neo-allotetraploids. Spider graphs reporting comparisons of the expression levels (in RPM) of the 21-nt (**B**) and 24-nt siRNAs (**C**) derived from *Copia* and *Gypsy* LTR-retrotransposons across the different libraries to the expression levels of siRNAs derived from LINEs and TIR transposons. They demonstrated the contrasting expression dynamics of siRNA populations according to the TEs the siRNAs originated from. *Gypsy*-derived 21-nt siRNAs were highly expressed in the S1 and S5 generations of the neo-allotetraploids while the other TE-siRNAs were mainly additively expressed. The *Gypsy* and *Copia*-derived 24-nt siRNAs were mostly expressed in the S5 generation, while the other TE-derived siRNAs were significantly under-expressed in the S1 generation. Annotation of TE-siRNAs was performed according to the TE classification published by Wicker et al. (2007).

Here, we have demonstrated the contribution of siRNAs derived from LTR-retrotransposons to the relative increase of the 21-nt class observed at the generation S1. However, less than 1% of the 21-nt siRNAs present in the RNA fraction of the S1 generation were annotated as LTR-retrotransposon-derived sRNAs. We thus pursued the characterization of other genomic components that may be responsible for the huge deviation in sRNA expression patterns observed in the S1 neo-allotetraploids.

### Specific populations of anonymous and virus-derived 21-nt siRNAs were increased in the S1 generation of the neo-allotetraploids

We filtered the remaining 21-nt siRNAs (not corresponding to miRNAs nor to TE-derived siRNAs according to our current annotation) on the basis of their overexpression in the S1 generation only. We obtained a total of sixty-eight siRNA sequences with expression levels ranging from 501 to 7192 raw read counts in the S1 generation, and only 0 to 6 reads in each of the parental and S5 generations (two examples are provided in fig. 6*A* and fig. 7*A*); they represented a total of 109 548 reads in the S1 generation, 7.23% of the 21-nt class (details in supplementary table S5, Supplementary Material on line). We assayed the relative abundance of a set of ten of these S1-specific 21-nt siRNAs by qRT-PCR. The initial F1 hybrids as well as the S0 and S3 generations of the three resynthesized *B. napus* allotetraploids were also included in the qRT-PCR experiments. In agreement with bioinformatic predictions, we demonstrated a significant increase of the siRNA reads in the S1 generation compared to the diploid progenitors and the S5 generation (with ΔCq>10 between the S1 generation and the diploid parents as well as between the S1 and S5 generations; supplementary fig. S3, Supplementary Material on line). Interestingly, qRT-PCR analysis of the F1 hybrids and S0 generation led to similar results as those obtained for the S1 generation, with a few cases where higher amounts of accumulation were depicted in S0, while results obtained from the S3 generation were closer to those generated from the S5 generation (fig. 6*B* and supplementary fig. S3, Supplementary Material on line).

**Figure 6.**
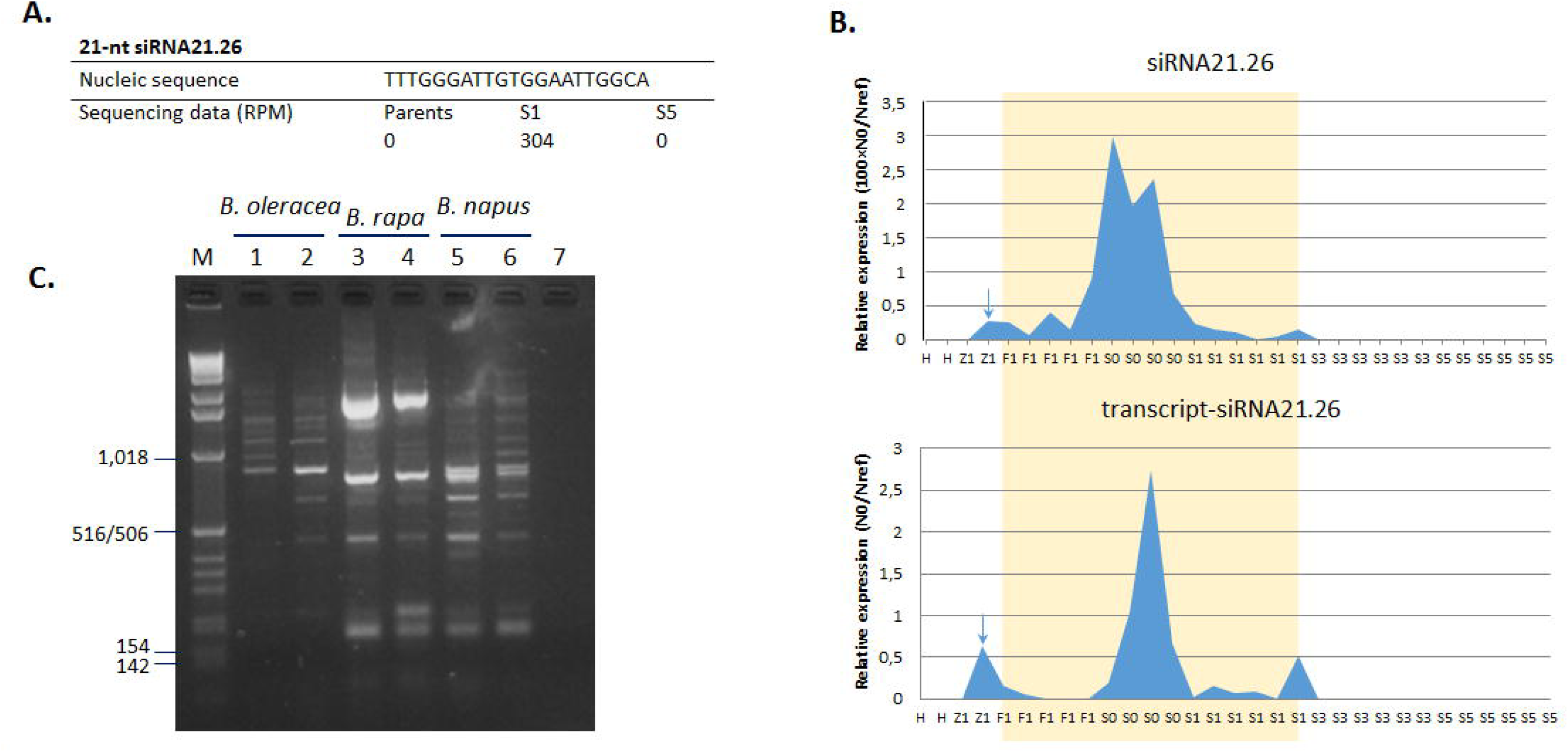
Unexpected virus-like transcripts appeared reactivated in response to allopolyploidy and subjected to regulation by specific siRNAs. (**A**) The 21nt-siRNA ‘siRNA21.26’ was part of the 21-nt siRNAs specifically accumulated in the S1 generation according to sequencing data. (**B**) Quantitative RT-PCR amplifications validated the accumulation of ‘siRNA21.26’ in the early generations of the neo-allotetraploids only, mainly in the S0 generation (orange area); they also demonstrated the same transcriptional dynamics of the transcript putatively targeted by ‘siRNA21.26’ corresponding to a virus-like sequence (with strong homology to #X13062). The arrows indicate unexpected contamination of cDNA (see Fig. S3 p.8/16 and p.14/16). (**C**) PCR reactions on *Brassica* DNAs using primers designed in the corresponding virus-like transcript (forward=5’-CGGGTTCCTCGTCTAC-3’; reverse=5’-CTCTACAAAGGCAATGGTT-3’) generated multi-band profiles, suggesting a dispersed location across the genome as truncated copies. *Legends:* diploid progenitors H, *B. oleracea* ‘HDEM’ and Z1, *B. rapa* ‘Z1’; F1, first hybrids; S0, doubled F1 hybrids; S1/S3/S5, selfing generations of the neo-allotetraploids; M: 1kb ladder; lanes 5 and 6: oilseed rape varieties Darmor and Yudal, respectively; lane 7: H_2_O as negative control.

**Figure 7.**
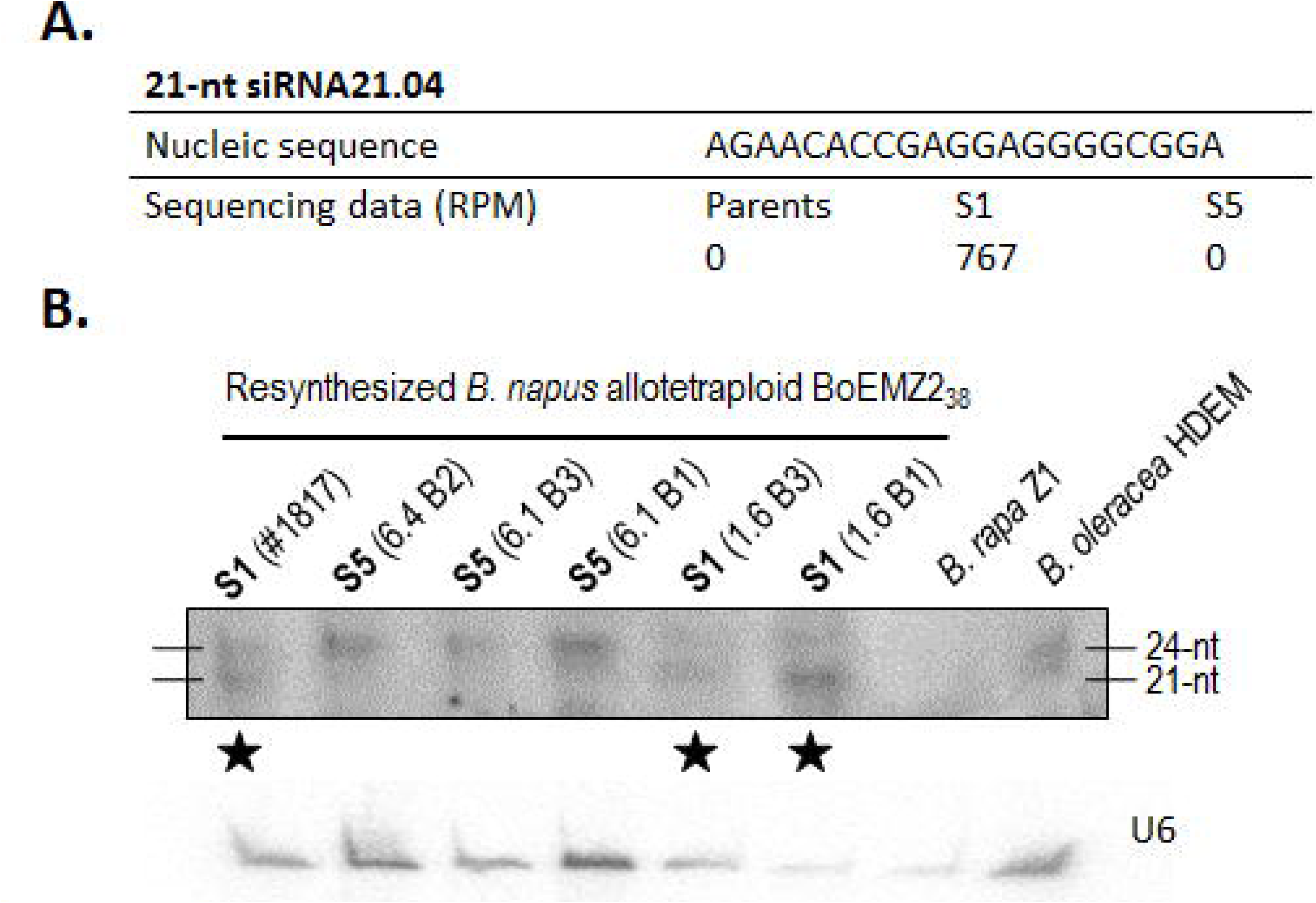
Validation of the differential expression levels estimated for the 21-nt small interfering RNA siRNA21.04. (**A**) Bioinformatic estimates from the analysis of sequencing data predicted high levels of expression in the S1 generation only (767 RPM against no reads in both the parental and S5 generations). (**B**) The expression levels were validated by Northern blot analysis with hybridization signals at the size of 21-nt only in the three different S1 genotypes analysed (⋆) (here from the neo-allotetraploid BoEMZ2_38_ for which expression at 527 RPM was estimated; see supplementary table S5). Additional signals at the size of 24-nt were identified in the two generations S1 and S5, with stronger signals in S5.

We undertook the identification of the molecular origins of these S1-specific 21-nt siRNA populations. No clear information was retrieved from exploitation of the *Brassica* genome sequences publically available. Nevertheless, exploitation of unannotated *B. napus* scaffolds available to us led to sequence alignments between 15 and 20-nt in length, with no mismatch, for all the sixty-eight siRNAs analysed, with a total of 246 different *B. napus* unannotated scaffolds retrieved (data not shown); the siRNAs under survey may thus originate from genomic sites that were not integrated in the published assembled *Brassica* genomes. We also exploited *B. napus* mRNAseq data; the data were generated from high-throughput sequencing of RNAs extracted from meiocytes of two different oilseed rape varieties that were both analysed at two different ploidy levels, haploid vs. diploid (NCBI BioProject PRJNA362706, Lloyd et al. 2018). Interestingly, twenty-nine of the sixty-eight 21-nt siRNAs to be characterized (43%) matched perfectly with mRNAseq reads that were present only in the haploid forms of one of the two varieties surveyed (list of the corresponding siRNAs provided in supplementary table S5, Supplementary Material on line). In particular, the two 21-nt siRNAs ‘siRNA21.26’ and ‘siRNA21.39’ matched respectively with seven and three reads that could be assembled in 319bp and 210bp-long sequences; by sequence similarity search with BLAST, the assembled expressed sequences showed high similarities with plant viruses (E-values of respectively 1e-165 with 99% identity covering 319bp of sequence #X13062, and 4e-76 with 92% identity covering 210bp of sequence #HQ388350). The 319bp-long expressed sequence also showed high similarity with a *B. napus* EST, previously identified when analyzing the population of transcripts in young flowers and flower buds (E-value of 3e-90 with 96% identity covering 201bp of the cDNA sequence #CX187929). Four of the retrieved mRNAseq reads matching four different siRNAs of interest (the virus-like derived ‘siRNA21.26’ and ‘siRNA21.39’, in addition to ‘siRNA21.21’ and ‘siRNA21.56’) were quantified by qRT-PCR; they displayed the same dynamics of expression as the corresponding siRNAs: the results revealed significant transcript amounts in the F1 hybrids, the S0 and S1 generations, but no expression in the diploid progenitors or in the S3 and S5 generations (fig. 6*B* and supplementary fig. S3, Supplementary Material on line). PCR amplifications on *Brassica* DNA extracts, using primers designed in each of the two viral-like transcripts of interest, gave multiband profiles (fig. 6*C*) suggesting a dispersed location of these putative virus-derived sequences in the *Brassica* genome; this recalls the main features of endogenous viral elements, EVEs, that were demonstrated to be present in all eukaryotic genomes (Feschotte and Gilbert, 2012), and mainly disseminated across the genome as truncated copies (Bertsch et al. 2009).

Altogether, our data have demonstrated the high mobilization of specific 21-nt-siRNA populations in the very first generations of the neo-allopolyploid formation, including the early step of the interspecific hybridization represented by the F1. Their mobilization was concomitant with the expression of non-coding transcripts that also appeared specific to the early generations of the resynthesized *B. napus* allotetraploids.

### Production of 21-and 24-nt siRNAs similar in sequence was highly reorganized during the early stages of the neo-allotetraploid formation

Expression levels were assayed by Northern blot hybridization for four of the sixty-eight 21-nt siRNAs that displayed significantly higher levels of accumulation in the S1 generation; they were considered as anonymous as their genomic origins could not be determined precisely. The S1-specific expression of the selected 21-nt siRNAs was clearly validated for three of them (namely ‘siRNA21.01’, ‘siRNA21.04 shown in fig. 7 and siRNA21.12). It was striking to observe additional 24-nt hybridization signals in both the S1 and S5 generations; the coexistence of populations of 21-and 24-nt siRNAs with similar nucleic sequences, and thus putatively involved in the silencing of the same targets, was observed in the S1 generation only (fig. 7*B*). Similarly, *in silico* mapping of the full sets of the sRNA reads, excluding miRNAs, on the two retrieved virus-like sequences corresponding to ‘siRNA21.26’ and ‘siRNA21.39’ (see above) indicated an over-representation of both 21-and 24-nt siRNAs similar to the viral sequences, in the S1 generation only (supplementary fig. S4, Supplementary Material on line). Characterizing the fraction of 21-nt siRNAs corresponding to TEs or anonymous sequences that mapped perfectly (no mismatch) onto 24-nt reads demonstrated the concomitant accumulation of 21-and 24-nt siRNAs similar in sequence as being particularly marked in S1, with more than 60% of the 21-nt siRNAs having 24-nt counterparts, against ~30% for the parents and S5. Expression dynamics of the corresponding 24-nt siRNAs was independently established by qRT-PCR experiments for two independent 21-nt/24-nt pairs (supplementary fig. S3, Supplementary Material on line). The patterns of sRNA accumulation thus indicate that distinct populations of siRNAs, involved in two different regulation pathways, coexist, have the same targets and appear to be specifically reorganized in response to allopolyploidy (fig. 8).

**Figure 8.**
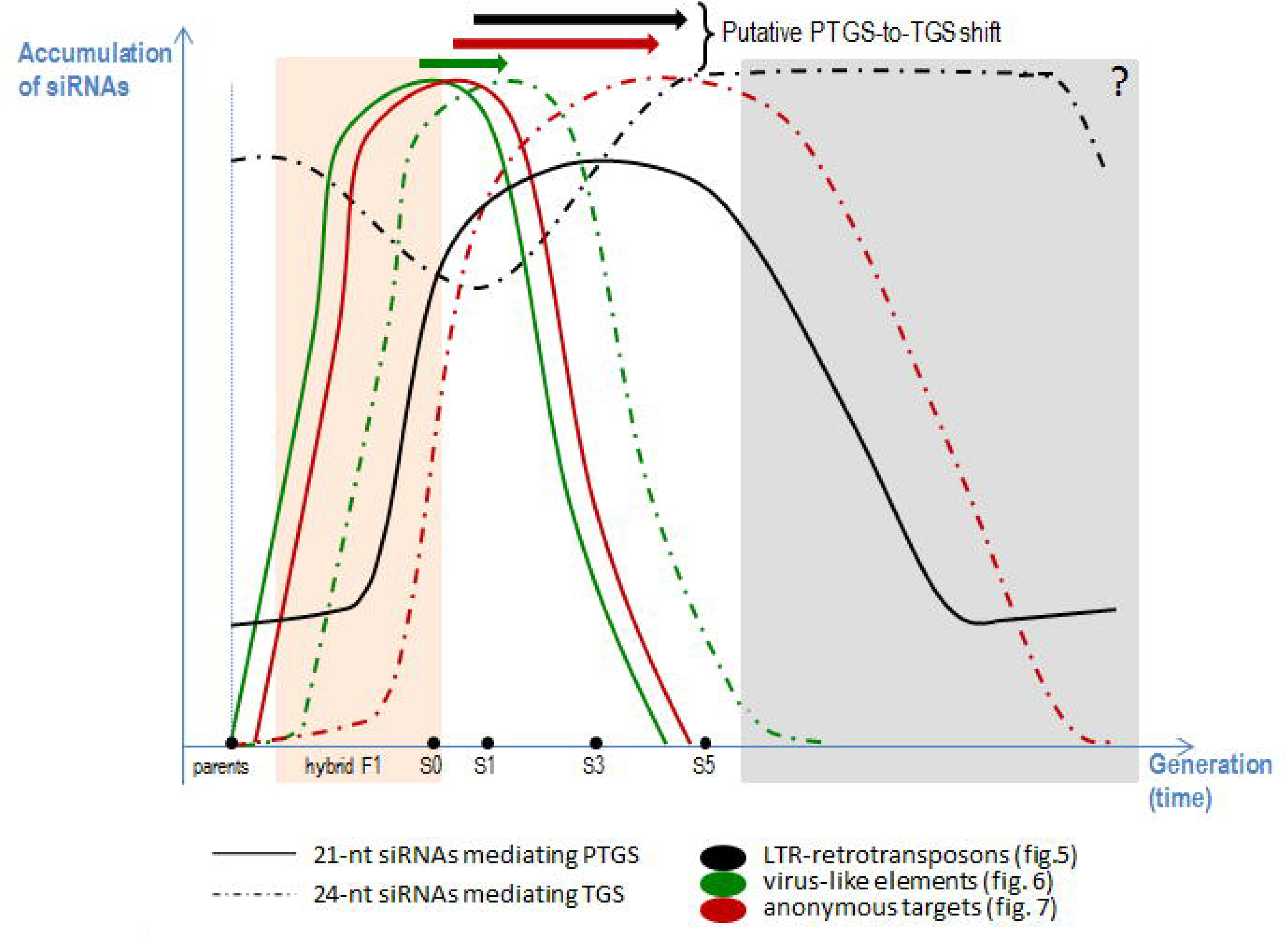
The global dynamics of siRNAs in response to allopolyploidy is characterized by the coexistence of two main silencing pathways leading to the progressive shift from PTGS mediated by 21-nt siRNAs to TGS mediated by 24-nt siRNAs during the formation of the allopolyploid genome. The global dynamics of siRNAs along the allopolyploidization process represented here is based on the results reported in the present study and summarized in fig. 5, fig. 6, fig. 7 and supplementary fig. S4. While the grey portion of the figure is hypothetical, the orange part corresponds to the interspecific hybrids F1 as well as the generation S0 that displayed significant levels of expression for siRNAs highly expressed in the S1 generation. We can thus wonder if the mobilization of the PTGS siRNA pathway is not the “emergency” genomic response to the genomic shock engendered by the interspecific hybridization step itself. The TGS pathway would then represent the longer established silencing pathway that ensures the stability of the neo-allopolyploid genome.

## Discussion

The present study reveals non-stochastic regulation of specific sRNA populations in response to allopolyploidization. Contrary to previous studies (Ha et al. 2009; Kenan-Eichler et al. 2011), our experimental design used “allopolyploid replicates”, enabling us to demonstrate that variations in sRNA accumulation were reproducible in three independently resynthesized *B. napus* allotetraploids. The immediate and coordinated changes in sRNA expression observed may suggest an immediate but necessary remobilization of sRNAs to face the genomic shock that accompanies the early stages of allopolyploidization. Annotation and identification of the sRNA populations reorganized during allopolyploidization and the characterization of their accumulation dynamics add to our knowledge of the major regulatory processes required to ensure the emergence of a new allopolyploid genome.

### Small RNAs experience immediate and transient accumulation changes in response to allopolyploidy

Our study demonstrates that small RNA populations experienced rapid and transient changes in the early generations of *Brassica* neo-allotetraploids. Most notably, quantitative analyses revealed that the 21-nt sRNAs were over-represented while the 24-nt sRNAs were under-represented, in the S1 neo-allotetraploids, in comparison with the relative accumulation levels estimated for both the parental and the S5 generations. Of course, the variability in the 21 and 24-nt fractions between the parental replicates (see details in table S1) might influence the MPV used as the reference, but statistics were performed to keep into account such variations, focusing on the significant differences across generations only. Additional experiments by qRT-PCR have validated such accumulation dynamic patterns, supporting the differential mobilization of sRNAs from different size classes in response to allopolyploidization.

Annotation of the 21-nt sRNA fraction indicated global miRNA stability across the three different generations. Interestingly, this relative miRNA stability echoes a previous study in hybrid and allopolyploid wheats that depicted global additivity of miRNA profiles in comparison with the progenitors (Kenan-Eichler et al. 2011). Despite the global additivity, both our analysis, and that undertaken in wheat (Kenan-Eichler et al. 2011), identified specific miRNAs that deviated from the expected MPVs in early allopolyploid generations. Interestingly there was some overlap in these non-additive miRNAs, e.g. miR168, and these represent interesting candidates to be further investigated when considering the regulation of specific functional targets in response to allopolyploidy. In contrast to the observations in wheat and *Brassica*, numerous deviations from MPVs were identified for the miRNA fraction in Arabidopsis allopolyploids, i.e. *A. suecica* and neo-allopolyploids (Ha et al. 2009). These contrasting patterns for the mobilization of miRNAs in response to allopolyploidy between different polyploid models thus indicate specific and more complex mobilization of miRNAs following allopolyploidy than generally expected.

The detailed annotation of the 24-nt sRNA fraction across the three different generations indicated a decrease in the relative abundance of the 24-nt TE-derived siRNAs in the S1 *Brassica* neo-allotetraploids consistent with previous studies on allopolyploids or hybrids (*Arabidopsis suecica*:Ha et al. 2009; wheat:Kenan-Eichler et al. 2011; rice:He et al. 2010, *Arabidopsis thaliana*:Groszmann et al. 2011, maize:Barber et al. 2012). In these studies, the decrease in 24-nt siRNAs corresponding to TEs and other repetitive sequences occurred concomitantly with the high TE transcription activity usually observed in the first generations of neo-allopolyploids (wheat:Kashkush et al. 2003, Kenan-Eichler et al. 2011; Arabidopsis:Madlung et al. 2005). Interestingly, the detailed annotation of TE-derived siRNAs we conducted identified different dynamics of TE regulation depending on their classification, with differential accumulations of 21-and 24-nt TE-derived siRNAs across the generations. The specific case of *Gypsy* retrotransposons can be underlined. We observed that *Gypsys* were the preferential TE targets of siRNAs following allopolyploidy, and that might be related to their preferential clustering in pericentromeric regions compared to other LTR-retrotransposons such as *Copias* mainly shown to be dispersed along the chromosomes (Alix et al. 2005). Reorganization of the neo-allopolyploid genome would particularly affect heterochromatic regions, with changes in epigenetic marks relaxing the epigenetic control of the concerned TEs like *Gypsys*, thus leading to the production of new siRNAs preventing further transposition events. In a previous study performed on the same *Brassica* plant material (Sarilar et al. 2013), we established that allopolyploidy had a moderate impact on restructuring at insertion sites for different TEs, and we identified copies of the *Athila*-like retrotransposon (clustered in pericentromeres) as being particularly quiescent. Our current data reinforce the hypothesis that specific siRNA-mediated TE repression mechanisms set in immediately after allopolyploidy to avoid anarchic transposition across the genome.

Nevertheless, with respect to the TE-specificity of the repression mechanisms identified in the present study (targeting mainly LTR-retrotransposons, the major TE fraction of plant genomes), some copies of specific TEs may escape this control (i.e. we identified transposition events for the transposon *Bot1* in *Brassica* neo-allotetraploids; Sarilar et al. 2013). The stability and fate of the neo-allopolyploid genome would depend on the efficiency of the small RNA pathways to control the populations of relaxed TEs inherited from both progenitors.

### Small RNAs might contribute to the success of allopolyploids

Analysis of small RNA populations targeting protein-coding genes clearly demonstrated that proteins harboring changes in their abundance immediately after allopolyploidisation (Albertin et al. 2007) were the preferential targets of sRNA-based regulation. Such results support our previous hypothesis that small RNAs might play a major role in the regulation of gene expression in response to allopolyploidy (Marmagne et al. 2010). We notably identified a significant over-representation of genes related to stress responses and metabolism (based on biological process GO terms), within the genes specifically targeted by non-additively accumulated sRNAs. The majority of these sRNAs displayed a significant decrease in their accumulation levels in the neo-allopolyploids compared to the diploid progenitors, indicating that the corresponding genes might have their transcription deregulated. Nevertheless, such deregulation of gene transcription seems to be quite specific, as a similar GO term enrichment analysis revealed that stress-related genes did not deviate significantly from additivity when considered in their global functional category (Albertin et al. 2007). Altogether, our results may thus indicate that the dynamics of sRNAs in response to allopolyploidy could contribute to the reprogramming of some specific genes involved in response to stresses as well as in plant metabolism and thereby provide support to the “genome shock” hypothesis (McClintock 1984).

The lower-accumulation of sRNAs involved in metabolism is in accordance with previous works performed on the same *Brassica* material that identified up-regulation of genes involved in metabolism and notably photosynthesis, in early allotetraploids (Marmagne et al. 2010) as well as on other *Brassica* neo-allotetraploids with diverse metabolic processes that were enriched in up-regulated genes in the first generation of selfing (Wu et al. 2018). Other studies dealing with the outcome of allopolyploidy have demonstrated the higher photosynthetic capacity of allopolyploids compared to their diploid progenitors (e.g. Arabidopsis, Ni et al. 2009, Solhaug et al. 2016; soybean, Coate et al. 2012). Here, our results indicate that small non-coding RNA pathways might have a role in the regulation of metabolic genes at the origins of the transgressive phenotypes of allopolyploids in photosynthesis and biomass production (i.e. growth vigor) that have contributed undoubtedly to their success over their diploid counterparts.

### Severe changes observed in the relative amounts of siRNAs indicate a PTGS-to-TGS shift at the first stages of neo-allopolyploid formation

One of the striking features of our results was the delayed accumulation of 24-nt siRNA populations compared to the accumulation of 21-nt siRNA populations, with both populations involved in the regulation of the same targets, during formation of the *Brassica* neo-allopolyploids. These dynamics are reminiscent of the shift from PTGS mediated by 21-nt siRNAs to TGS mediated by 24-nt siRNAs (Mirouze 2012) that was demonstrated to accompany *de novo* silencing of the active and transcribed plant LTR-retrotransposon *EVD* in Arabidopsis (Marí-Ordóñez et al. 2013): *EVD* reactivated transcripts were first subject to the PTGS pathway with a high accumulation of 21-nt siRNAs, and then *EVD* transcript levels declined in response to TGS corresponding to an accumulation of 24-nt LTR-derived siRNAs, thus maintaining epigenetic silencing by RdDM of the active retrotransposon. As expected from Marí-Ordóñez et al. (2013), we observed coexistence of populations of 21-and 24-nt siRNAs with similar nucleic sequences in the S1 generation, with the former being progressively replaced by the latter. Interestingly, the speed of this shift appeared to be dependent on the targets repressed, i.e. immediate (in S1) for endovirus-like sequences but more progressive (from S1 to S5) for LTR-retrotransposons.

In addition, it is now established that the AGO1 protein has a crucial role in the antagonistic miRNA-and siRNA-mediated RNA silencing pathways through PTGS, with miR168 being demonstrated as the key regulator of AGO1 homeostasis by repressing *AGO1* mRNA and thus limiting the availability of the AGO1 protein for efficient siRNA-mediated PTGS (Mallory and Vaucheret, 2009; Martínez de Alba et al. 2011). Interestingly, miR168 is one of the few micro RNAs that displayed changes in accumulation across the generations, validated by qRT-PCR. Our data support the set-up of a highly efficient 21-nt siRNA-mediated post-transcriptional silencing in the very first generations of the neo-allopolyploid (S1), with low levels of miR168 and active AGO1, that is replaced in the following generations (S1 to S5) by the 24-nt siRNA-mediated transcriptional gene silencing, with increased miR168 accumulation and AGO1 repression. In the polyploid wheat model, only over-representation of miR168 was described (Kenan-Eichler et al. 2011), suggesting that efficient 24-nt siRNA-mediated RNA silencing by TGS is an almost immediate response to allopolyploidy in the wheat system.

Altogether, our results support a hypothetical model whereby the response to allopolyploidization entailed a PTGS-to-TGS shift (fig. 8). It can be assumed that specific regions (enriched in LTR-retrotransposons, endovirus-like and other anonymous sequences) became transcriptionally active upon allopolyploidization due to the deregulation of RdDM-based TGS caused by genomic shock (with DNA methylation changes as usually observed in neo-polyploids; Hegarty et al. 2011 and references therein). This transcriptional reactivation then triggered immediate mobilization of 21-nt siRNAs produced from the transcribed sequences to enhance PTGS and jugulate the proliferation of these usually tightly repressed sequences. Finally, 24-nt siRNAs were produced to re-establish TGS, which progressively succeeded PTGS. During this transition, distinct 21-and 24-nt siRNA populations transiently coexisted in the neo-allopolyploid, with the specific generations of overlap determined by the speed with which the regulatory system could be (or had to be) set up.

The present study contributes to our knowledge of the coordinated regulation of small RNA pathways that optimize genome stability in response to the genome stress triggered by allopolyploidy. In this respect, small interfering RNAs may be considered guardians of genome integrity during the first steps of neo-allopolyploid formation (Malone and Hannon 2009). Accordingly, it can be now questioned whether the success of a new allopolyploid event might be related to this PTGS-to-TGS shift and its effectiveness, in the immediate and long-term, in controlling reactivated TEs and other non-coding sequences. This is particularly true given that these non-coding sequences appear to play a significant role in the outcome of interspecific crosses and allopolyploidization events (Martienssen 2010).

## Materials and Methods

### Plant material and RNA extraction

Our plant material consisted of three independently resynthesized *Brassica napus* allotetraploids (i.e. resulting from three independent crosses but involving the same diploid progenitors) namely BoEMZ1_38_, BoEMZ2_38_, BoEMZ5_38_ and their diploid progenitors *B. oleracea* (‘HDEM’, female donor) and *B. rapa* (‘Z1’). It comprised three different generations of neo-allopolyploids: S1, S3, S5 (fig. 1) from which the S1 and S5 generations were used for sRNA sequencing (see below). At least two cuttings per plant were grown under controlled conditions (18°C during 8h night and 21°C during 16h) in the same climate chamber, and samples were collected individually from each single cutting, all at once. The first hybrids F1, the corresponding doubled hybrids S0, and S1 individuals from one previous experiment conducted in 2004 (samples were stored at −80°C; Albertin et al. 2006) were integrated into the present study for molecular validation of specific small RNAs and transcripts. All plant material was produced by A.-M. Chèvre and collaborators, and has already been used in several studies in which it was described in more detail (*e.g.* Szadkowski et al. 2010; Sarilar et al. 2013). Total RNAs were extracted from stems using TRIzol^®^ following the manufacturer’s protocol (Invitrogen), with addition of 0.2 μl/ml beta-mercapto-ethanol to TRIzol^®^ extemporarily. We collected and used stem tissues in order to analyse the same plant tissue from the same plant material as previous studies focusing on protein expression changes during *Brassica* allopolyploidisation (Albertin et al. 2006, 2007) to allow integration of the data.

### Small RNA library construction and sequencing

Small RNA library construction and sequencing using the Illumina GA II sequencing system were performed by ImaGenes GmbH (Berlin, Germany). In total, eleven small RNA libraries were constructed and sequenced, corresponding to RNA extracts from stems of the diploid progenitors (with two biological replicates for each progenitor) as well as a 1:1 parental RNA mix used as a technical replicate, and RNA extracts from stems of the three resynthesized *B. napus* allotetraploids using the two generations of selfing S1 and S5 (fig. 1).

### Bioinformatic analyses

#### Quality control and pre-processing

A total of 23,102,628 sequence reads were obtained after Illumina sequencing of the eleven small RNA libraries that were specifically constructed for the present study (fig. 1). The quality control tool for high throughput sequence data FastQC (Version 0.10.1; http://www.bioinformatics.babraham.ac.uk/projects/fastqc/) was used for validation of the sequencing data to be further surveyed. Sequences of adaptors, as well as empty reads, low complexity reads and reads smaller than 18-nt and bigger than 26-nt were removed by using Filter Tool from the UEA sRNA toolkit (Moxon et al. 2008). In a subsequent step, Filter Tool was used to remove reads similar to transfer RNAs (tRNAs) and ribosomal RNAs (rRNAs). To increase the sensitivity of Filter Tool, we built *Brassica*-specific tRNA and rRNA databases from *de novo* predictions using locally the tRNAscan-SE 1.21 software (Lowe et Eddy 1997; Schattner et al. 2005) and the RNAmmer 1.2 software (Lagesen et al. 2007) on the genome sequence of *Brassica rapa* (*Brassica rapa* Genome Sequencing Project Consortium, Wang et al. 2011). We predicted a total of 1138 tRNA sequences and 79 rRNA sequences; these sequences were then added to the whole set of t/rRNAs registered in the Rfam 10.1 RNA family database (Griffiths-Jones et al. 2003). Sequence reads matching t/rRNAs (no mismatches allowed, 100% coverage) were eliminated from our present analysis. Finally, a total of 14,917,073 sequence reads were further analysed.

#### Quantification of sRNA populations and normalization

Each small RNA library was analyzed individually; read counts were normalized to allow direct comparisons between sRNA data from the different libraries (see below, statistical analysis). Size distribution of sRNAs was first computed for comparisons between generations. Then, full annotation of the sRNA classes of interest was performed to tentatively identify their origins.

#### Identification of known and putative microRNAs

Each of the eleven short-read libraries was searched for known miRNAs using miRProf (from the UEA sRNA toolkit, Moxon et al. 2008) that identifies sequence reads matching already registered miRNAs in miRBase (version 19, database: plant_mature, with the options: group-organisms, group-family and collapse-match-groups were enabled; Griffiths-Jones 2004). A locally developed Perl script was used to estimate expression levels for each known microRNA retrieved from our sequence data; results were summarized in supplementary table S3, Supplementary Material on line. In addition, we performed *de novo* predictions to identify putative miRNAs using miRCat (from the UEA sRNA toolkit, Moxon et al. 2008) with the following options, according to the documentation: --genomehits 16 --hit_dist 200 --maxgaps 3 --max_overlap_length 70 --max_percent_unpaired 60 --max_unique_hits 3 --maxsize 24 --min_abundance 5 --min_energy −25.0 --min_gc 10 --min_hairpin_len 75 --min_paired 17 --minsize 20 --percent_orientation 80 --pval 0.1 --window_length 100. Two different predictions were performed using either the *B. rapa* genome sequence or the *A. thaliana* sequence; results are provided in supplementary table S4, Supplementary Material on line (the results from miRCat with all the annotation details can be provided upon request). The quantification of small RNA was conducted as described above.

#### Identification of small RNAs derived from transposable elements

Sequences of small RNAs that were not identified as miRNAs were compared to the Repbase database and a home-made *Brassica* transposable element database (Sarilar V, Martinez Palacios P, Joets J, Alix K, unpublished data) using Blast (word size = 21 or 24), and only perfect matches were further considered. Annotation of individual TE-derived siRNAs was made according to the TE class / order / superfamily they originated from, following the TE classification from Wicker et al. (2007).

#### Search for the genomic origins of small RNAs of interest

Search for the genomic origins of the sRNA populations specifically expressed in the S1 generation was first performed by alignments of the reads against *Brassica* genome resources (genome of *B. rapa*, Wang et al. 2011; genome of *B. napus*, Chalhoub et al. 2014; GSS from *B. oleracea*, from J. Craig Venter Institute) using the Bowtie 2 software (Langmead and Salzberg 2012) with no or only one mismatch allowed (setting k=10). We exploited *B. napus* mRNAseq data that were generated from high throughput sequencing of RNAs extracted from meiocytes of two oilseed rape varieties originating from two different germplasms (namely Darmor and Yudal) and surveyed at two different ploidy levels, i.e. haploid (AC) and euploid (AACC) (NCBI BioProject PRJNA362706, Lloyd et al. 2018). The mRNAseq reads that contained an exact match to the small RNAs under survey were extracted from FASTQ files using a custom PERL script. After multiple sequence alignment of the retrieved mRNAseq paired reads using ClustalW2 (http://www.ebi.ac.uk/Tools/msa/clustalw2/), the resulting mRNA sequences were used as queries for similarity searches against the non-redundant nucleotide collection at NCBI (http://blast.ncbi.nlm.nih.gov/Blast.cgi) with the programme BLASTN.

The sRNA sequence data have been deposited in the Gene Expression Omnibus (GEO) database at NCBI; accession number GSE94076.

### Statistical analysis of differential sRNA expressions supported by NGS data

Among each library, read counts were normalized to adjust for differences in library size and coverage, allowing direct comparisons between sRNA data from different libraries; each raw read count was multiplied by 10^6^ and then divided by the total read count of the whole library, the unit becoming ‘reads per million’, RPM. When considering expression levels of individual sRNA sequences, only sRNAs with normalized expression levels ≥ 30 RPM within each generation were analysed, taking into consideration the work from Fahlgren et al. (2009) who demonstrated highly reproducible data for relatively abundant sRNAs with log_2_(RPM)>5. Analysing the patterns of size distributions of sRNAs for each library allowed the identification of an enrichment in the 18-20-nt classes for the library corresponding to the S5 BoEMZ1_38_ line (fig. 2*A*), suggesting a higher amount of RNA degradation that may generate some background noise when estimating sRNA expression levels; data originated from this library were thus discarded for further statistical analyses based on individual library sequencing data. To identify differences in expression for specific sRNA populations, we performed a one-way analysis of variance, ANOVA. The classification variable was the generation with five different levels *B. oleracea* ‘HDEM’, *B. rapa* ‘Z1’, 1:1 parental RNA extract mix, S1 and S5. As the differences between the profiles obtained from the 1:1 parental RNA extract mix and the mean of the two parents were not found significant (see below), we calculated the mean parent value (MPV) as the weighted mean of ‘HDEM’, ‘Z1’ and of the 1:1 RNA extract mix from the two diploid parents. We then performed the three following contrasts: MPV vs. S1, MPV vs. S5, and S1 vs. S5 (using a Student’s *t*-test procedure).

When estimating significant allopolyploidy-related differences in expression of sRNAs between the diploid progenitors and the different resynthesized *B. napus* allotetraploid lines, sRNA expression data in the neo-allopolyploid lines were compared to the mid-parent expression value (MPV). When analyzing size distribution of sRNAs across the generations, the calculated mid-parent value (MPV_c_), corresponding to the average expression data estimated from the four individual parental sRNA libraries, was compared to sRNA data obtained for the library constructed from the 1:1 parental RNA mix which represented a technical replicate (MPV_t_). As no significant differences were found (with *P-*values such as 0.18≤*P*≤0.96), the MPV taken as the reference for all the comparisons we made corresponded to the mean value of sRNA expression data obtained from the five parental libraries.

### Analysis of the relationships between the dynamics of sRNAs and protein expression patterns

A previous comparative proteomics study was performed on the same *Brassica* material as the one used in the present work, in order to identify proteins specifically affected by allopolyploidy (Albertin et al. 2007). From a total of 377 stem proteins, we focused our analysis on a set of 263 proteins (not taking into account proteins with several isoforms) that included 152 proteins depicted as non-additive proteins and 111 as additive proteins. Each protein was characterized by its function and one *Arabidopsis thaliana* protein-coding gene (i.e. Atxgxxxxx). Each *A. thaliana* gene was used to retrieve all the *Brassica* homologous genes from the two diploid species *B. rapa* and *B. oleracea* to constitute ‘*Brassica* functional group sequences’ (one group per protein): we exploited published data from the analysis of the complete *B. napus* genome sequence (Table S19 from Chalhoub et al. 2014) and complemented the search for homologues with BRAD (http://brassicadb.org/brad/) using BLAST against the *B. rapa* and *B. oleracea* genome sequences. Genomic (including the 500bp upstream and downstream regions) and cDNA sequences were retrieved from the database ‘EnsemblPlants’ (http://plants.ensembl.org/index.html) using Biomart. Using Galaxy from GQE – Le Moulon (Moulaxy, http://galaxy.moulon.inra.fr) and for each of the eleven sequenced libraries, we mapped the whole populations of 21-and 24-nt sRNAs separately, on the genomic and the cDNA sequences in parallel; we used Blast (word size = 21 or 24) and only perfect matches (100% similarity) were further considered. Redundancy was carefully removed (filtering using *unique line remove duplicate lines*, not to count one specific read more than once) when resuming mapping results for one ‘*Brassica* functional group’; we finally obtained the total number of reads targeting a specific function, for each of the 11 libraries, allowing differentiation between putative genomic or transcript targets. After normalization, statistics were performed as described above, comparing sRNA expression data in the neo-allopolyploid to MPV. We further analysed sRNA data for ‘*Brassica* functional groups’ with at least 1 RPM in one library and ddlR≥1 (from ANOVA). GO term annotation (focusing on biological processes) and enrichment analysis of the protein-coding genes putatively targeted by the non-additive sRNAs (*P*≤0.01) was performed using GO::TermFinder with default parameters (Boyle et al. 2004); we visualized the results using REVIGO (Supek et al. 2011).

### Real time quantitative reverse transcription PCR (qRT-PCR) analysis

RNA extracts were treated with DNAse I (DNA-*Free*Ambion #1906). First-strand cDNA was synthesized from 500 ng of total RNA using the miScript II RT kit (Qiagen), using the miScript HiFlex Buffer in order to allow quantification of both sRNAs and mRNAs. Real-time quantitative RT-PCR (qRT-PCR) for sRNAs was performed using miScript SYBR Green PCR kit (Qiagen) on a 7500 Real-Time PCR System (Applied Biosystems). Reactions were performed following manufacturer’s instructions, by adding to the mix 0.5 μM of the oligonucleotide corresponding to the direct sequence of the targeted sRNA and 5 μl of the synthesized cDNA diluted 300 times, and subjected to the following programme: 95°C for 15 min, 40 cycles of 94°C for 15 s, Ta for 30 s and 70°C for 45 s; Ta ranged from 55°C to 65°C and was optimized for each primer. Quantitative RT-PCR assays were performed on the diploid parents, the three F1 hybrids and the corresponding S0 doubled hybrids, and the following generations of selfing S1, S3, S5; two cuttings were tested (except for two F1 hybrids and two S0 lines for which only one plant was available), and at least two technical replicates were performed.

The qRT-PCR data were analysed using the LinRegPCR software v.2012.x (Ruijter et al. 2009) to determine the PCR efficiency for each sample. The mean PCR efficiency per reaction and the Cq value were then used to calculate a starting concentration N0 per sample. Relative expression levels were obtained from the ratio between the N0 of the sRNA under survey and the normalization factor Nref, N0/Nref (with Nref corresponding to the mean N0 value of the additively expressed micro RNAs miR159a, miR160a, miR166a, which were first verified for their highly stable expression in our samples using GeNORM (Vandesompele et al. 2002), as shown in supplementary fig. S1 (Supplementary Material on line)). Following the same procedure and on the same cDNA templates, we also assayed transcript amounts for transcripts retrieved by similarity search against *B. napus* meiosis mRNAseq data. The different primers used for both sRNA and mRNA qRT-PCR experiments are provided in supplementary fig. S1, fig. S2 and fig. S3 (Supplementary Material on line).

Only qRT-PCR data points for sRNAs and mRNAs with significant expression levels (Cq≤35) were retained and N0/Nref ratio were calculated and used for further statistical analyses, when required. The ΔCt method using miR159a and miR160a as references was applied to the analysis of the dynamics of accumulation of miR168. Significant expression changes across generations were estimated using a one-way ANOVA then completed by pairwise comparisons using Student’s *t*-test with no adjustment.

### Small RNA blot analysis

Small RNA blot analysis with 15 μg of RNA per sample was performed as described in Lelandais-Brière et al. (2009) using [γ-^32^P]ATP end-labeled oligonucleotide probes complementary to the sRNA sequences to be analysed, with an oligonucleotide probe complementary to U6 snRNA used to normalize RNA concentrations. Hybridization was performed at 50°C overnight using labeled probes at ~5×10^6^ cpm/membrane; 2 washes with 5×SSC 1% SDS and a third wash with 2×SSC 1% SDS at 50°C for 10 min each were performed; pictures were taken 5h after exposure at −80°C.

## Acknowledgements

We particularly thank Clémentine Vitte (CNRS, GQE-Le Moulon, Gif-sur-Yvette, France), Jérôme Gouzy (INRA, UMR LIPM, Toulouse, France) and Mark Tepfer (INRA, IJPB, Versailles, France) for valuable discussions. We thank Pierre Montalent (INRA, GQE-Le Moulon, Gif-sur-Yvette, France) for technical assistance with Galaxy. Three anonymous reviewers are also acknowledged for their helpful comments on the manuscript. PMP was supported by a PhD fellowship from the French *Ministère de l’Enseignement Supérieur et de la Recherche* (MESR). Experiments dedicated to small RNA library construction and sequencing were funded by the *Institut Fédératif de Recherche 87 ‘La plante et son environnement’* (Gif-sur-Yvette, France) and the *Direction Scientifique* of AgroParisTech (Paris, France). This paper is dedicated to the memory of our colleague Dr Hervé Thiellement.

